# EMT associated transcription factors Slug and Snail demonstrate specific expression patterns in breast cancer subtypes, involving tumor or stroma components and correlating with proliferation status and complex clinical prognosis

**DOI:** 10.64898/2026.05.29.728692

**Authors:** Flora Doffe, Lucile Mercier, Damien Drubay, Benjamin Verret, Natasha Joyon, Pierre Savagner

## Abstract

EMT associated transcription factors (EMTaTF) controls the epithelial-mesenchymal transition (EMT) process, but their detailed expression pattern in cancer is poorly documented at the cellular level. Here, we located two major EMTaTF: Snail and Slug by immunochemistry using validated antibodies in a cohort of 569 invasive breast carcinomas. We screened all tumor molecular types using TMA analysis in addition to full tumor sections, identifying cell types and structures involved in their expression, correlated to morphological, functional, and clinical characteristics. Briefly, Slug was expressed in Luminal A/B and HER2 breast cancer subtypes by almost all cells from the basal layer in normal-looking tubule structures and was very sporadically seen in transformed cells in the *in situ* or invasive components. In contrast, Slug was visibly expressed in 28% of our triple-negative (TN) tumor samples, significantly expressed by invasive tumor cells. Slug was also strongly expressed in a large subpopulation of stroma fibroblast-like cells in all breast carcinoma subtypes. Conversely, Snail was frequently expressed in transformed cells in all invasive breast carcinomas subtypes, up to 54 and 64% of tumor cells in TN and HER2 tumors respectively. Expression pattern was heterogeneous and included invasive areas and in situ component. Some stroma cells, particularly endothelial cells from the tumor microenvironment were also found to express Snail. Slug and Snail proteins were also detected in metastatic foci, sometimes with an increase in the expression level, particularly for Snail. Unexpectedly, we found a significant positive correlation between cell proliferation and Slug stroma cell expression in luminal A and B subtypes. This link bolstered the relative proximity we uncovered between CK- KI67- Slug+ stoma cells and CK8+ KI67+ Slug- tumor cells in LumB samples. However, survival long-term studies failed to demonstrate a link between slug tumor or stromal expression and long-term survival. On the other hand, Snail protein expression in tumor cells was surprisingly and significantly linked to a better survival fate overall. In contrast, Snail overexpression in stroma cells from primary tumors was significantly linked to a time-dependent tendency for relapses, metastasis and poorer survival, arising when post-surgery time lapse increased. In conclusion, these findings offer new and unsuspected understanding of the complex localization pattern and clinical involvement of Snail genes in breast cancer, beyond classic EMT pathways.

## Introduction

The expansion of resistant tumour cell clones during cancer treatment is a critical issue in the clinical management of most carcinoma. This emergence reflects the adjustment of tumor microenvironment, promoting heterogeneity, immune resistance and inducing cell subclones that express dedifferentiated phenotypes. These phenotypes reflect partially their original epithelial cell type, but also the reactivation of oncogenic and developmental pathways silenced during embryogenesis. Among those, epithelial-mesenchymal transition (EMT) pathways have been well described to control cell differentiation and fate during development, inducing individualization and motility. During this process, so-called EMT associated transcription factors (EMTaTF) control cell organization and phenotype, including cell-cell and cell-matrix structures. Importance of the full EMT process during tumor progression is debated in the cancer field, but EMTaTF can clearly impact clinically relevant cell responses (Lüönd et al., 2021; Thompson et al., 2025; Yang et al., 2020; Ye et al., 2017) including immune response^3^. Surprisingly, their precise expression pattern in tumors remains poorly documented. In addition, their topological link with tumor areas supporting proliferation and invasion was not precisely established in clinical samples.

We focused in this study on breast cancer, characterizing the expression pattern of two members of the Snail family: Slug and Snail, in all subtypes. Previous studies have reported meta-analysis approaches linking the expression of EMTaTF with breast cancer progression (Imani et al., 2016). These reports, based on RNA or protein expression levels establish a significant correlation between the expression of EMTaTF and unfavorable outcome. However, they do not make a distinction between breast cancer molecular subtypes, and they do not take in consideration the cancerous, epithelial or stroma cell types actually expressing EMTaTF, limiting their clinical significance. Overall, Snail and Twist family was usually described to be the most involved in the clinical progression Snail (Snai1) and Slug (Snai2) genes were described in several studies, including ours, to play a role in breast tumor progression (Nieto et al., 2002, Come et al., 2006). Slug was usually described, as a protein, to be linked to unfavorable clinical outcome in terms of OS and sometimes Distant Free Survival (DFS) (Zhang et al., 2022), but again, these studies did not identify the cell populations expressing Slug and their respective contribution.

Slug gene is expressed during development and causally involved in several EMT stages, including gastrulation and neural crest cell migration (Nieto et al., 1994). It is expressed in adults in most pluristratified epithelia and in the breast in basal/myoepithelial cells (Nassour et al., 2012). It is clearly involved in mammary epithelial cell basal differentiation and fating, providing progenitor/stem cell properties, proliferation potential, motility and participating to tubulogenesis (Guo et al., 2022; Nassour et al., 2012; Phillips et al., 2014), in link with P-cadherin as a direct target and effector (Idoux-Gillet et al., 2018). Slug involvement in tumor initiation has been demonstrated in a MMTV-PyMT mouse model comparable to the tumor-initiating capabilities of Epcam-positive tumor cells. Using the same mouse model and live-markers, Ye et al. (Ye et al., 2015)described Slug and Snail distinct expression pattern along the tumor progression, linking Snail to luminal-derived breast cancer cells, when Slug was restricted to the basal lineage, as described previously. However, The experimental inhibition of Snail expression in breast cancer cell line MDA-MB-231 cells resulted in a notable increase in their ability to invade in vitro and in vivo in mice engrafted with these cells, suggesting more complex mechanisms at play involving FOXA1 upregulation in this case (Tsirigoti et al., 2022). Overall, a clear need appears for more documented studies describing Slug and Snail expression pattern in a large clinical cohort. In this manuscript, we exploited a 569 well characterized invasive breast carcinomas cohort.

## MATERIALS ET METHODS

### TMA

A large scale cohort was initiated in 2011 in Gustave Roussy in collaboration with Sanofi. It was elaborated to include all breast cancer molecular subtypes and only patients with a standard of care procedure including first line surgery and second line treatments to only get tumor tissue intact with at least a 5-year follow-up and complete clinical database. This TMA is available to clinicians and researchers internally and externally after assessment of the project by the ethical committee and project leaders. This cohort called GRAND_TMA includes 1800 patients, and includes tissues fixed with A.F.A (Alcohol, Formaldehyde, Acetic acid) and Formaldehyde. To avoid variation in staining due to those two fixative methods we filtered on the Formaldehyde fixative reagent that is now on the only used, resulting in a subcohort of 569 patients.

### EMTaTF antibodies validation

Slug commercial antibody (C19G7 Rabbit mAb #9585, Cell Signaling Technology) was validated by several laboratories, involving Slug-KO mouse control in our work (Nassour et al, 2012) for immunocytochemistry. Snail antibody (AF 3639 Goat BioLegends) was also validated by several laboratories for immunocytochemistry (Gonnissen et al., 2017). We found it to provide a very similar localization pattern to the monoclonal anti-Snail antibody produced, validated and published by AG De Herreros’s laboratory (De Herreros et al., 2010). To support the relevance of our findings, we only considered nuclear staining for Slug and Snail, more specific and easier to use for quantification.

### Colorimetric staining

The histological colorations are done on Ventana discovery Ultra (Roche) using Slug Rabbit mAb), with an anti-rabbit polymer HRP revealed by DAB (3,3’-Diaminobenzidine) chromogen. Snail Goat antibody was used adding an enhancer (kit Polink 2 plus anti-goat) and an anti-goat polymer (kit Polink 2 plus anti-goat), revealed with DAB chromogen. A counterstaining (Hemalun de Mayer) was used to mark all nuclei. We scanned the slides with Nanozoomer Hamamatsu. Histoscore (H-score) was calculated as a semi-quantitative assessment of both the intensity of staining and the percentage of positive cells. Counting and analysis were performed using Qupath 0.3.0. to create a TMA de-arrayer and detect positive cell staining and sorting analysis between stroma and cancer area using subtracting masks

### Multiplex immunofluorescence

The histological colorations are conducted on Ventana discovery Ultra (Roche) using the antibodies obtained from several sources: Slug (C19G7, rabbit, Cell signaling C19G7 #9585), Keratin 5 (PRB160P, Poly19055, BioLegend, San Diego, CA, USA), Keratin 8 (904804, MMS-162P, BioLegend), Ki-67 (MIB-1, M7240, Unconjugated, Agilent Dako Roche), Hoechst (33342, Sigma Aldrich). The secondaries antibodies are either anti-rabbit Discovery UltraMap anti-Rb HRP (Roche 760-4315) or anti-mouse Discovery UltraMap anti-Ms HRP (Roche 760-4313) and the tertiaries antibodies anti-HRP are Opale 520 (FP1487001KT, Akoya), Opale 570 (FP1488001KT), Opale 620 (FP1495001KT, Akoya), and opale 690 (FP1497001KT, Akoya). The tissue slides are imaged at all fluorescence lasers (450, 488, 555, 675 nm) with the Vectra microscope (automated quantitative pathology imaging system, Perkin Elmer). The software Inform (Akoya) unmixes the fluorochromes. This last software allowed analysis of fluorescent positive cells and segmentation depending on the area (cancer/stroma). To analyse the cohort of patients, each with more than 50 images, we have used the R plugin PhenopTr on R for Windows v4.2.2 and its package Rtools 4.2 (4.2.0.1). We have used the same plugins to analyse the distances between each phenotype.

### Clinical endpoints

The prognostic endpoints considered were the invasive disease-free survival (iDFS), the distant disease free-survival (DDFS), and the overall survival (OS). The iDFS was defined as the time from surgery to the first occurrence of any recurrence, new cancer, or death from any cause. The DDFS was defined as the time from surgery to the first occurrence of distant recurrence or death from any cause. The OS was defined as the time from surgery to death.

### Statistical methods

Several statistical tests were used, based on the data format, including unpaired T test, Welch T test and 2 way ANova. H-score values were first normalized by square root transformation. All other analyses were realized using the R software (v 4.2.1). Descriptive statistics were computed with the *crosstable* package. Spearman correlations and their p-values were computed with the *cor.test* function. The survival analyses were performed using the *survival* package. The continuous variables presenting distribution with heavy tail (Snail/slug expressions, tumor size, positive nodes, and Ki67) were normalized with the Yeo-Jonhson transformation (from the *bestNormalize* package) to limit the influence of extreme values. The survival analyses were conducted with the multivariable Cox model, adjusted for the available classical risk factors (age at surgery, tumor size, tumor grade, luminal status, number of positive nodes, and the Ki67). The proportional hazard assumption was checked with the Schoenfeld residuals trend test and visual representation. The time-transform function *tt()* was used to consider an interaction between the biomarker and the time when the proportional hazard assumption did not hold.

## RESULTS

### Study population

From the original subcohort, we removed the patients without information about the stromal/tumoral Slug/Snail expressions or the clinical risk factors used in this study (age at surgery, tumor size, tumor grade, luminal status, Her2 status, number of positive nodes, and the Ki67), resulting in a final dataset of 569 patients. The descriptive statistics of these patients are presented in the Table 1.

**Table 1:** Breast carcinoma cohort description.

| label | variable | value |
| --- | --- | --- |
| Age at surgery | Med [IQR] | 59.1 [49.8;70.4] |
|  | Min / Max | 27.2 / 94.8 |
| Tumor size | Med [IQR] | 21.0 [15.0;30.0] |
|  | Min / Max | 5.0 / 112.0 |
| Tumor grade | 1 | 96 (16.87%) |
|  | 2 | 271 (47.63%) |
|  | 3 | 202 (35.50%) |
| Number of positive nodes | Med [IQR] | 0 [0;2.0] |
|  | Min / Max | 0 / 44.0 |
| Luminal status | Luminal A | 317 (55.71%) |
|  | Luminal B | 164 (28.82%) |
|  | RE negative | 88 (15.47%) |
| Ki67 (%) | Med [IQR] | 11.7 [5.0;20.0] |
|  | Min / Max | 0 / 81.7 |
| Stromal Slug | Med [IQR] | 5.1 [1.9;11.5] |
|  | Min / Max | 0 / 58.8 |
| Tumoral Slug | Med [IQR] | 5.4 [2.0;10.2] |
|  | Min / Max | 0.1 / 108.3 |
| Stromal Snail | Med [IQR] | 0.1 [0.03;0.4] |
|  | Min / Max | 0 / 7.2 |
| Tumoral Snail | Med [IQR] | 2.4 [0.9;5.0] |
|  | Min / Max | 0 / 110.2 |

### Slug and Snail are expressed by distinct cell compartments among breast cancer subtypes

Aiming at a large-scale survey of breast cancer samples, we screened 18 tissue-macroarrays (TMA) slides displaying a total of 569 invasive breast carcinoma samples: 52 basal-like, 31 HER2+, 317 Luminal A (LumA), 164 Luminal B/ HER2+/- (LumB) tumors and 5 less common subtypes (Fig.1A). This cohort follows typical clinical recruitment range at Gustave Roussy. Three distinct representative core microsections were spotted on one slide from each tumor and analyzed independently using QuPath software. Considering the historic difficulty identifying reliable antibodies for Slug and Snail, we first validated antibodies as described in the Material and Methods section. We first looked at the general Hscore in each subtype to find a strong heterogeneity, even within each tumor subtype, but showing no significant differences (Fig. 1B). We then scored separately tumor cells and non-transformed differentiated myoepithelial cells. This scoring clarified that myoepithelial cells were mostly responsible for Slug expression in the tumor area in luminal tumors. Conversely, TN and HER2 tumors that do not usually include normal-looking tubules did express Slug in invasive tumor cells at a significant level (28 and 11% respectively, Fig. 1C). Upon direct observation, we found that Slug was physiologically expressed in about 10-20% basal/myoepithelial cells in normal healthy mammary gland, corroborating our previous observations in mouse mammary lobes (Fig. 2 and Nassour et al. 2012). This percentage could increase up to 60-80% basal/myoepithelial cells in normal lobules close to tumor cell aggregates in some patients (Fig.1D). We scored Slug expression pattern in breast tumors by distinguishing 3 distinct cell type subpopulations: a) basal/myoepithelial cells from normal-looking mammary ducts located in the tumor area, potentially including tumor in situ component, b) invasive ductal carcinoma (IDC) clusters and c) stroma cells. We found Slug to be strongly expressed in most basal/myoepithelial cells in Luminal A/B and HER2 subtypes in apparently normal ducts, sometimes encasing *in situ* tumor cells that were Slug-negative (Fig. 2, 3). About 100% basal/myoepithelial cells expressed Slug, a much higher proportion than in normal ducts. This unmistakable pattern was very reminiscent of the expression pattern of P63, a classic basal/myoepithelial marker (Wang et al. 2002). In TN carcinoma, known to not include any ductal structures, we found Slug to be expressed by a subpopulation of invasive dispersed carcinoma cells in about 28% of our samples (Fig. 1C and 2). The percentage of Slug-expressing cells in these samples ranged from 5 to 50% of tumor cells. Slug+ cells were distributed apparently randomly among Slug- tumor cells in most cases. Slug was very rarely seen in large cohorts of invasive tumor cells from other tumor subtypes.

**Figure 1.**
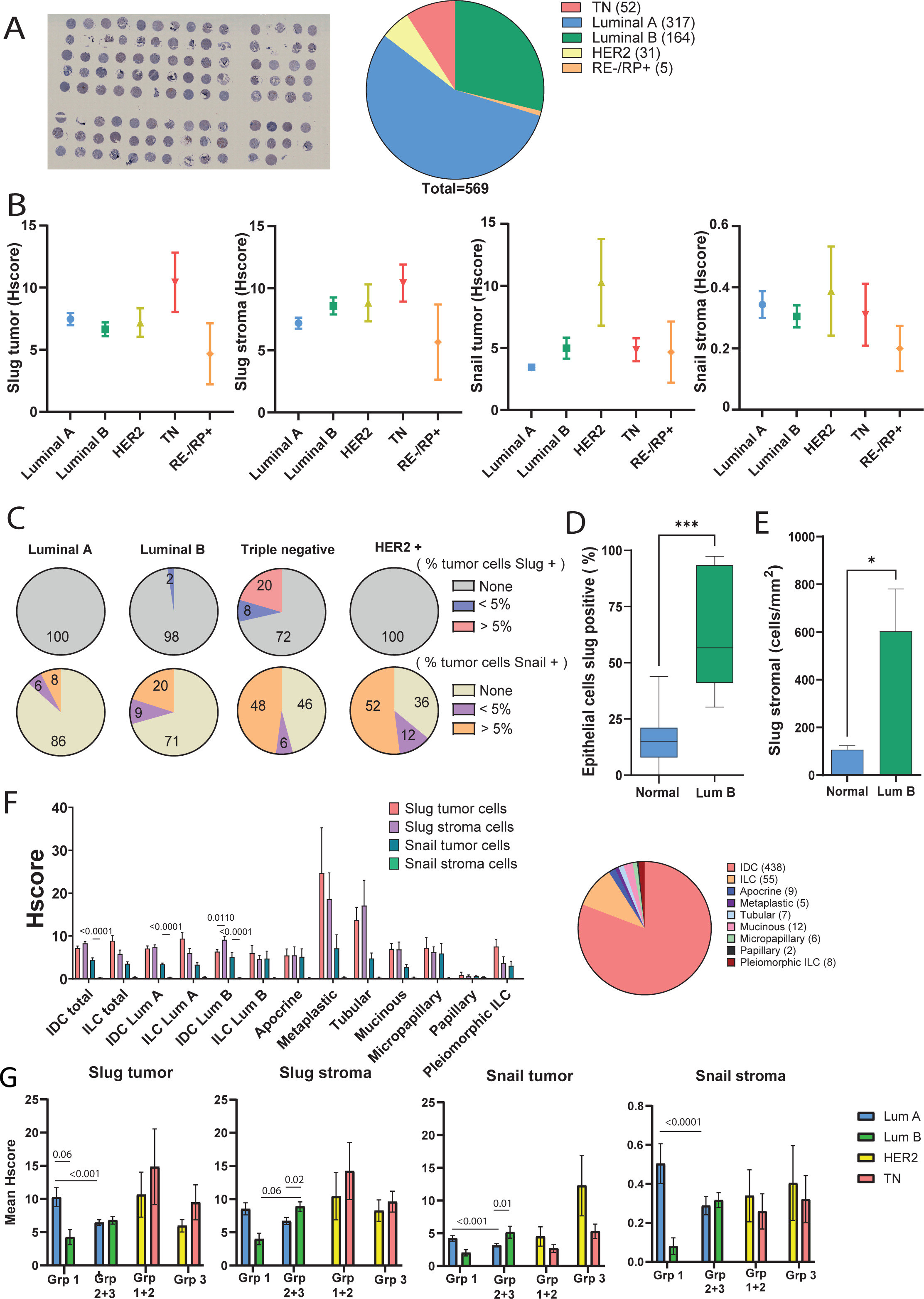
Slug and Snail expression pattern in breast carcinoma. A. Tissue-macroarrays (TMA) slides displaying a total of 569 invasive breast carcinoma samples included most breast cancer subtypes. B. Hscores were calculated from tumor or stroma cells for Slug and Snail protein expression pattern, demonstrating cancer subtype specificity. C. The percentage of tumors expressing Slug or Snail is displayed with a color-coded typing according to the percentage of tumor cells expressing Slug or Snail within each tumor. Normal-looking Slug+ myoepithelial cells were not included in these calculations. D. The percentage of normal-looking Slug+ myoepithelial cells was compared between 2 normal breast samples (NB: 12 fields of view) and 2 Luminal carcinoma samples (Lum: 14 fields of view), analyzed using QuPath software and single IHC Slug labeling. E. The percentage of Slug+ stroma cells was compared between normal breast and breast carcinoma samples based on multiplex analysis of 4 normal breast samples (16 fields) and 4 LumB carcinoma samples (19 fields) analyzed with inForm software (Akoya). F. Breast carcinoma histological subtypes express varying amount of Slug and Snail protein in tumor and stroma cells, showing significant differences in several cases. G. Slug and Snail protein expression level (Hscore) was also significantly linked to pathological grades in several instances. Unpaired T test two tailed was used for D with p=0.033 (*), p=0.002 (**) and p <0.001 (***), Welch T test was used for E, p<0.05 (*) and two-way ANOVA was used for F and G.

**Figure 2.**
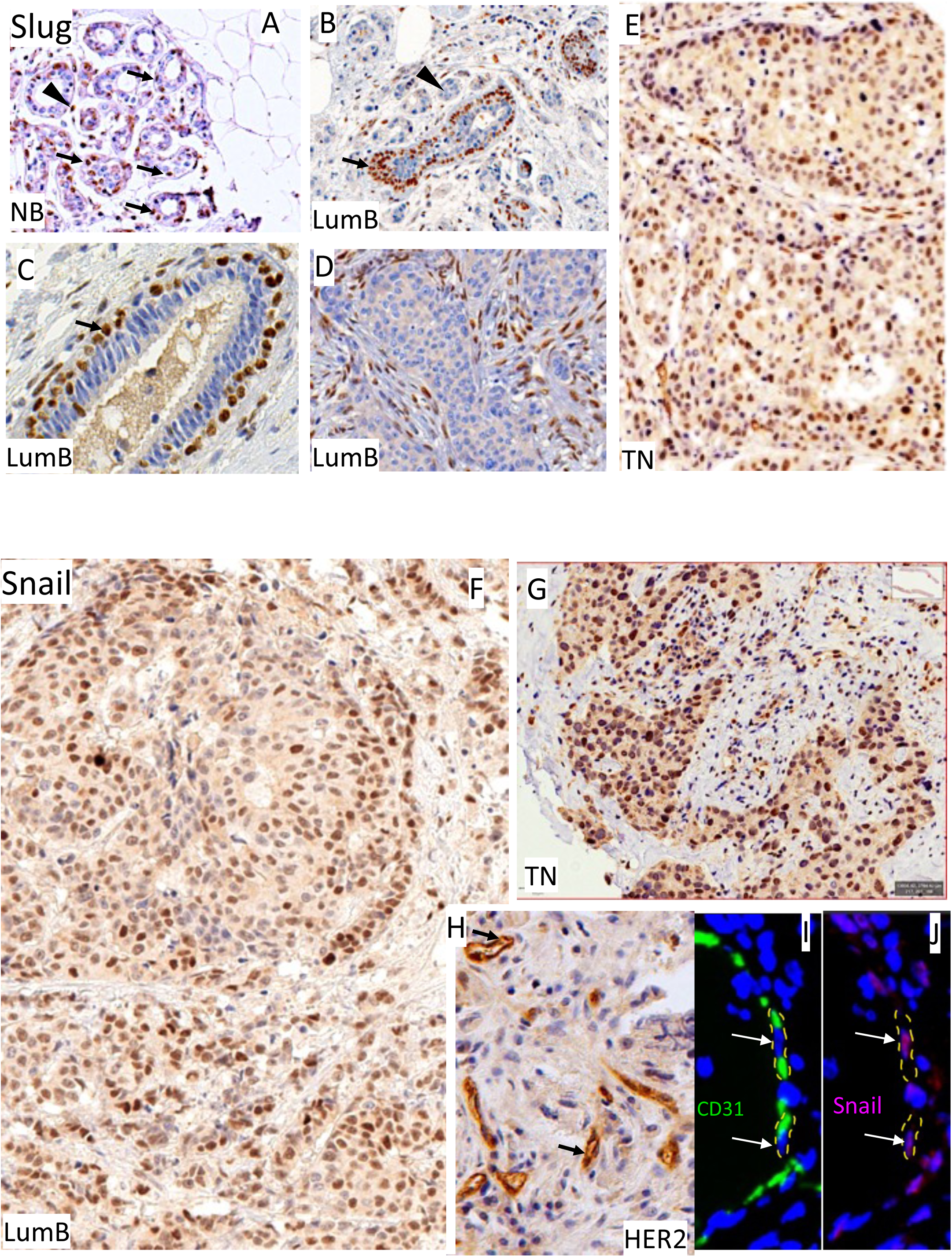
Localization of Slug and Snail in normal breast (NB) and breast carcinoma. A. Slug is seen in few basal myoepithelial cells around a tubule and in some stroma cells. B. Slug is expressed in most basal/myoepithelial cells delineating a tubule including in situ component within a LumB carcinoma. C. Higher magnification shows the Slug+ basal/myoepithelial cell organization. D. A significant percentage of fibroblast-like stroma cells express Slug. E. Slug is expressed in a varying percentage of invasive tumor cells in a TN breast carcinoma. F. Snail is found in invasive tumor cells in a LumB carcinoma. G. Snail displays a similar pattern in TN breast carcinoma. H. Snail is expressed in most visible endothelial cells in a HER2 breast carcinoma. I-J. Endothelial cells are double labeled for Snail (red) and CD31 (green) in the same tumor sample.

Stroma cells were found to express Slug significantly in all breast tumor subtypes. In normal healthy breast, this expression was limited (below 10% in peritubular region), but the expression level and the percentage of Slug-expressing fibroblasts strongly increased in tumor stroma, involving up to four-times more stroma cells in two thirds of our samples (Fig. 1E). Slug+ stroma cells were particularly abundant around Slug- invasive clusters invading adipose tissue (Fig. 3). Finally, a small percentage (< 10%) of the endothelial cells that we could identify by the HES staining were found to express Slug.

**Figure 3.**
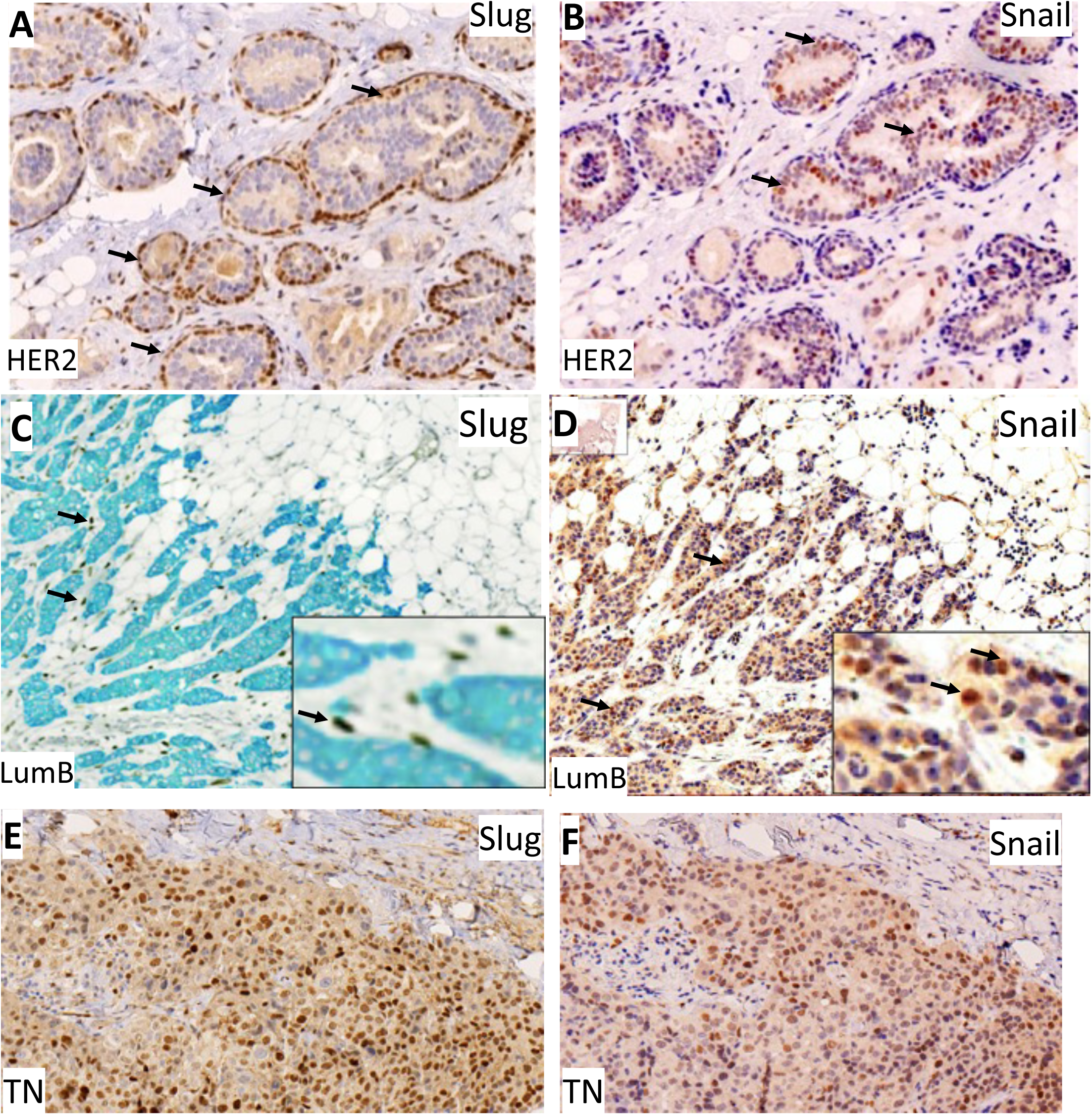
Slug and Snail expression patterns. AB. Two close slices of one HER2 breast carcinoma were labeled for Slug (A) or Snail (B). Arrows point to positive cells in the myoepithelial layer (A: Slug) or in the in-situ component (B: Snail). C. LumB breast carcinoma invasive front, double labeled for cytokeratins (blue) and Slug (black). D. A slice from the same paraffin block labeled for Snail show invasive tumor cell cohorts expressing Snail, as better seen in higher magnification inserts (lower right). E-F. Two close slices show tubules enclosing in situ tumor component, with Slug (E) and Snail (F).

Snail expression pattern was very different. We detected no Snail in normal breast tissue, beyond rare mesenchymal cells. In contrast, Snail was detected in disseminated cancer cells in invasive clusters from all subtypes, particularly in HER2 (64%) and TN tumors (64%) (Figs. 1-3). It was also sometimes expressed by cancer cells from the *in-situ* component in LumA (14%)/B (29%) tumors. Snail was typically expressed in invasive cohorts of tumor cells invading adipocytes (Figs. 2, 3). Much less Snail was seen in tumor stroma cells, as compared to Slug (Fig. 1). However, a strong expression level was found in most endothelial cells in the tumor area, not detected in normal breast (Fig. 2).

Slug and Snail were very rarely found to be co-expressed in the same tumor area. We recorded exceptional instances, only in TN carcinomas (Fig. 3. E, F).

### Stroma Slug expression correlate with proliferation marker in tumor cells in Luminal tumors

Using the same quantitation method, we analyzed our TMA set for the expression of the proliferation marker KI67. As expected, we observed that LumA tumor cells expressed less KI67 than other tumor subtypes. After calculating both Spearman and Pearson correlation coefficients, we unveiled an unsuspected correlation between the KI67 H-score in tumor cells and the Slug expression H-score in stroma cells in the LumA and B subtypes (Table 2). We also found a tendency in TN and HER2 tumors, when using the Spearman index. However, the smaller number of samples and probably the heterogeneity among tumors reduced the significance using the Pearson indexes.

**Table 2:** Spearman and Pearson correlation coefficient and associated risk between KI67 H-score in tumor cells and the Slug and Snail expression H-score in tumor and stroma cells.

|  |  | Spearman CC | p | Pearson CC | p |
| --- | --- | --- | --- | --- | --- |
| **Lum A**<br>**N=317** | Slug Tum | -0.004 | ns | 0.092 | ns |
|  | Slug Str | 0.188 | **< 0.001** | 0.220 | **< 0.001** |
|  | Snail Tum | 0.020 | ns | 0.008 | ns |
|  | Snail Str | 0.005 | ns | 0.040 | ns |
| **Lum B**<br>**N=164** | Slug Tum | -0.021 | ns | 0.129 | ns |
|  | Slug Str | 0.220 | **< 0.005** | 0.289 | **< 0.01** |
|  | Snail Tum | 0.014 | ns | 0.075 | ns |
|  | Snail Str | 0.175 | < 0.05 | 0.124 | ns |
| **Her2**<br>**N=31** | Slug Tum | 0.024 | ns | 0.073 | ns |
|  | Slug Str | 0.425 | < 0.05 | 0.258 | ns |
|  | Snail Tum | 0.433 | < 0.05 | 0.272 | ns |
|  | Snail Str | -0.078 | ns | 0.115 | ns |
| **TN**<br>**N=52** | Slug Tum | -0.049 | ns | 0.093 | ns |
|  | Slug Str | 0.282 | < 0.05 | 0.240 | ns |
|  | Snail Tum | 0.019 | ns | 0.040 | ns |
|  | Snail Str | 0.026 | ns | -0.055 | ns |

### Slug and Snail expression levels fluctuate with tumor histological subtypes

Among all breast cancer histological subtypes included in our cohort, tubular and metaplastic histological subtypes were found to overexpress Slug, mostly in basal/myoepithelial cell fraction for the tubular subtype, and in a mix of basal/myoepithelial and tumor cells for the metaplastic subtype (Fig.1F). In contrast, the lowest Slug expression level was detected in papillary subtype that express a more differentiated phenotype.

Snail expression was particularly strong in tumor cells from micropapillary and metaplastic breast tumor subtypes.

### Slug and Snail expression levels in tumor and stroma cells are linked to histology grades

In order to evaluate a potential link with tumor progression, we compared Slug expression levels (H-score) between distinct pathology grades established by Gustave Roussy pathology department, recording separately epithelial tumor/myoepithelial cells and stroma cells for all subtypes (Fig. 1G). In spite of the expected strong variability, we uncovered some statistically significant differences. We found that Slug expression level (H-score) in basal/myoepithelial cells from Grade 1 LumA tumors was about twice higher than the level observed in advanced grades 2 or 3, in a statistically very significant index (p< 0,001). We also observed a higher Slug expression in LumA, as compared to LumB grade 1 tumors, with a near significant value (p = 0,06). Snail was also more expressed in LumA tumor or stroma cells from grade 1 tumors, as compared to more advanced grades.

In addition, Slug stroma cell expression level in grades 2/3 LumB tumors was about twice higher than in grade 1, with a near-significant associated risk (p=0,06) in a unilateral test justified by the strong overexpression in higher-grade tumors and suggesting a link between stromal Slug and tumor aggressiveness in these highly proliferative tumors. It was also significantly lower in low-proliferating LumA stroma cells.

For HER2 and TN subtypes, we regrouped Grades 1 and 2 for because of the small number of grade 1 samples in our cohort. These groups showed stronger heterogeneity and we could not identify statistically significant differences.

### Slug and Snail can be modulated during the metastasis process

We compared Slug and Snail expression pattern in a group of ten primary breast cancer samples (LumA, LumB and TN) and in derived metastatic foci. Overall, Slug expression was not enhanced in Lum tumor-derived metastatic foci. However, it was induced in metastasis samples from our two Slug- primary TN samples. Snail induction or overexpression was observed in most metastatic foci derived from Snail- or Snail+ primary samples, (Fig. 4. CD).

**Figure 4.**
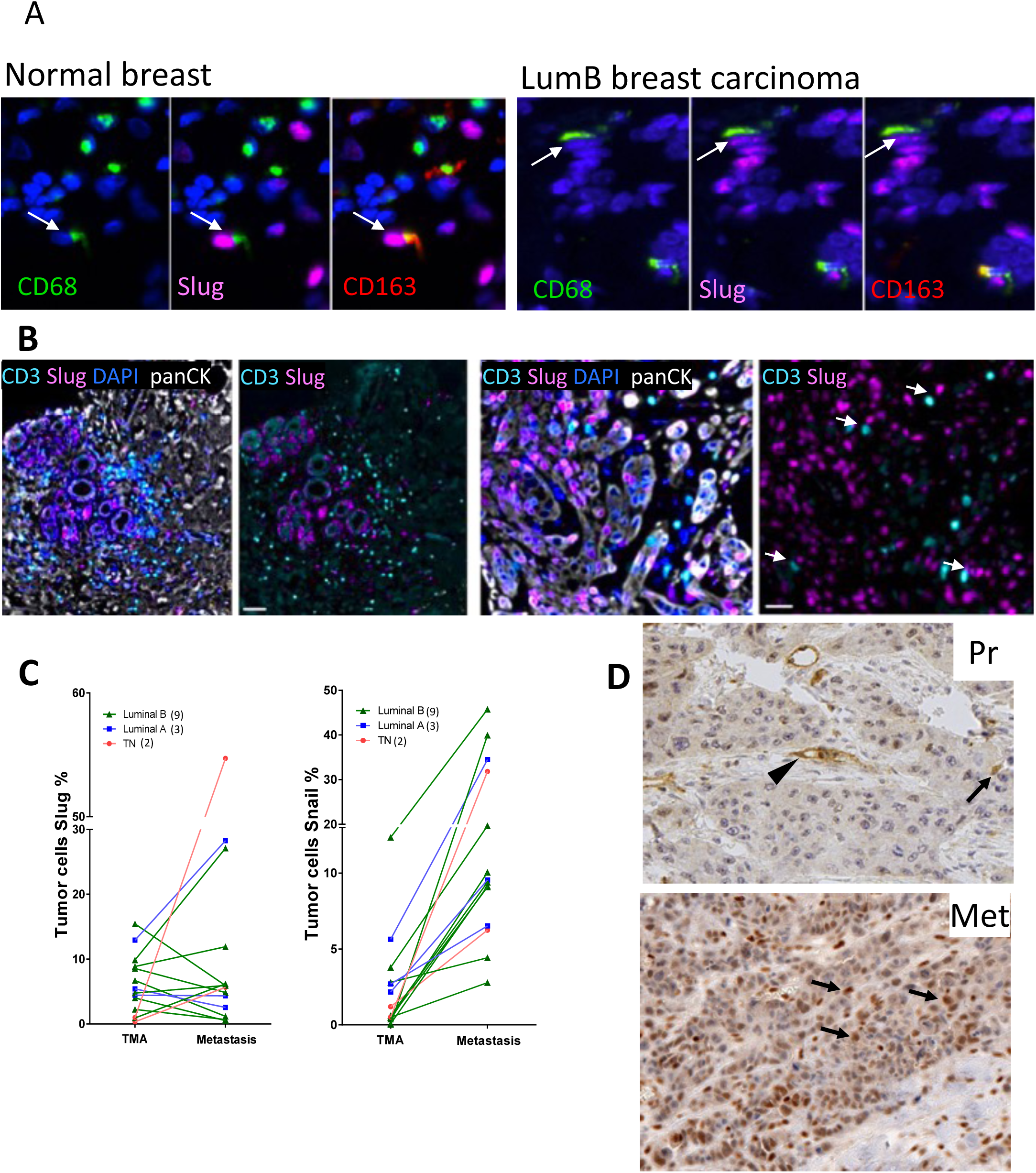
A. Slug is expressed in normal and tumor microenvironment macrophages. Few Slug+ cells are found to co-express Mac1 or Mac2 (arrows). B. Slug is not found to be co-expressed in CD3+ lymphocytes (arrows) in two LumB breast carcinoma samples with high (left panels) and low (right panels) lymphocyte infiltration. C-D. Slug and Snail expression in metastatic sites. Slug is not induced during the metastatic process, except in few cases. In contrast, Snail was induced or upregulated in most of our samples. D. Snail is upregulated in tumor cells (arrows) from a metastatic site (Met) derived from a primary tumor (Pr) expressing Snail only in non-tumoral stroma (arrow) and endothelial cells (arrowhead).

### Tumor multiplex analysis of Slug expression pattern in luminal B subtype

To understand on a larger topological scale Slug expression pattern in tumors, we analysed full-size tumor sections covering 300-600 mm^2^ from 3 Luminal B untreated tumor samples and performed double labelling for Pan-cytokeratin (PanCK) and Slug (Fig. 3C) Cells expressing Slug and cytokeratins (Slug+ CK+) were the myoepithelial-like cells we observed on TMA spots. Cells expressing Slug, but no cytokeratins (Slug+CK-) were the stroma cells also observed in TMA LumB spots. We focused on LumB tumors to take advantage of the restricted pattern of Slug expression in myoepithelial-like cells that we observed in this tumor subtype, easily distinguished from the stroma cells using image analysis. After quantification using Q-path software, we found that about 10% of CK+ cells were co-expressing Slug (corresponding to myoepithelial-like cells), when about 85% were CK+Slug-, presumably tumor cells. These results supported our TMA results.

To identify other Slug+ cell types, we performed fluorescent co-labelling of specific markers for fibroblasts, macrophages and lymphocytes. Overall, we found that a 15% stromal sub-population was expressing Slug. We also located cells expressing fibroblast activation protein (FAP), an activation marker for fibroblasts. After image analysis, we characterized several stroma subpopulations defined by their phenotype: 15% stroma cells were FAP+ Slug- cells, 10% were FAP- Slug+ cells and 5% were FAP+ Slug+ cells, emphasizing that about a third of the activated fibroblasts (FAP+) were expressing Slug. Other unidentified stroma cells also expressed Slug, presumably belonging to the immune component (Fig. 4. AB). We could not find CD3+Slug+ lymphocytes, but located rare CD68+ and CD163+ Slug+ macrophages.

We then developed a multiplex staining including Slug, CK5 (basal/myoepithelial cell marker), CK8 (luminal cells marker), and Ki67 for cell proliferation status to better understand cell dynamics and Slug expression pattern in the tumor (Fig.5). Using QuPath software, we could analyze spatial organization of cell subpopulations expressing our markers, separately in the stromal and tumor areas. Accordingly with our TMA study, we found that most luminal cells (CK8+) were Slug- negative and comprised within the tumor body (about 95%), within intact tubules defining the *in-situ* component, or migrating as cohesive clusters (Fig. 5H, 6C). Overall, tumor cells expressing Slug or Snail were found to display clearly epithelial features such as cytokeratin (5 or 8) and cohesiveness. In the rare cases of full tumor dispersion seen locally or on a bigger scale (invasive lobular carcinomas), no expression of Slug or Snail was noticed in isolated tumor cells (Fig. 5F). In fact, we could not see any induction of Slug in CK8+ during the tubule dissociation preluding to carcinoma invasion, when invading CK8+ cells were highly dissociated (Fig. 5F, Fig. 6A) or remained strongly cohesive (Fig. 5G). Slug was still expressed in CK5 cells during and after myoepithelial cell extensive dissociation and tubule breakup, evoking an EMT-like event Fig. 5B). We frequently noticed a limited CK5+CK8+ subpopulation of tumor cells suggesting some differentiation impairment (Figs. 5DH-6CD) involving about 3% of tumor cells (Fig.5H), typically in Slug- invading clusters. Remarkably, we mostly found KI67 expressed in Slug-negative cells (Fig. 5. C-G), a significant difference with the developing mammary gland where Slug is mostly expressed in a CK5+ proliferating cell compartment (Nassour et al. 2012). Snail antibody was found to be unfit for multiplex staining protocol, but classic IHC staining showed a noticeable Snail accumulation in cancer cells breaking up from tubules (Fig. 6B).

**Figure 5.**
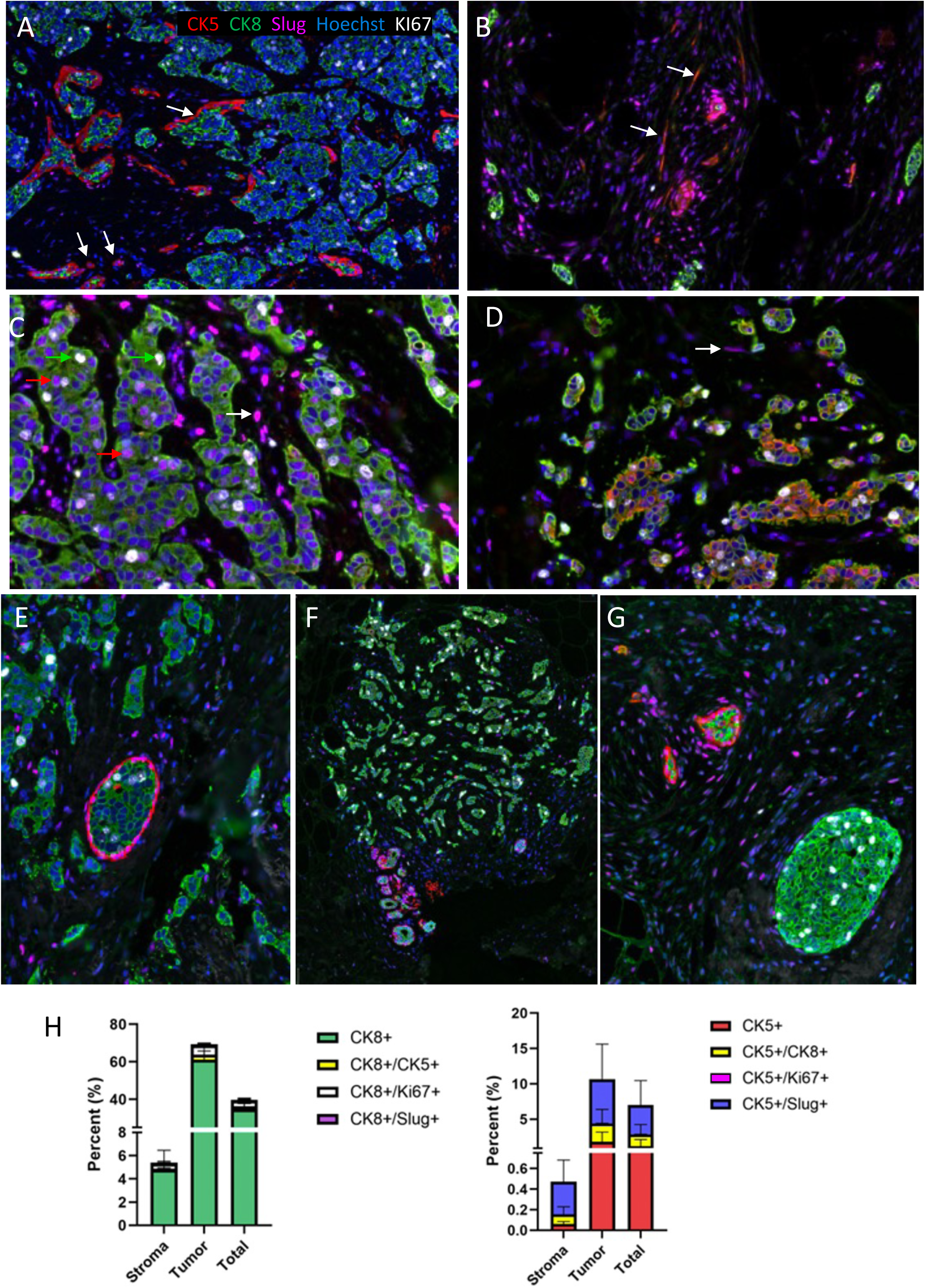
Multiplex analysis shows CK5, CK8, Slug, and KI67 expression pattern in Hoechst dye-stained tumor cells, within a LumB breast carcinoma. A-B. CK5+ myoepithelial cells (arrows) are strongly dissociated along mammary duct rupture and carcinoma invasion. C-D. Slug is mostly seen in stroma cells (white arrows) and CK5+ cells but rarely in CK8+ cells. KI67 is mostly expressed by CK8+ cells (green arrows). D. CK5 and CK8 can be coexpressed in tumor cells. E-G. CK8+ carcinoma cells leaving the tubules typically maintain some cohesiveness (E, G), but can be strongly dissociated (F). Quantification using QuPath software confirm these patterns: Most KI67+ cells are CK8+. Most Slug+ cells are stromal or CK5+. A small but significant percentage of CK5+ cells are also CK8+.

**Figure 6.**
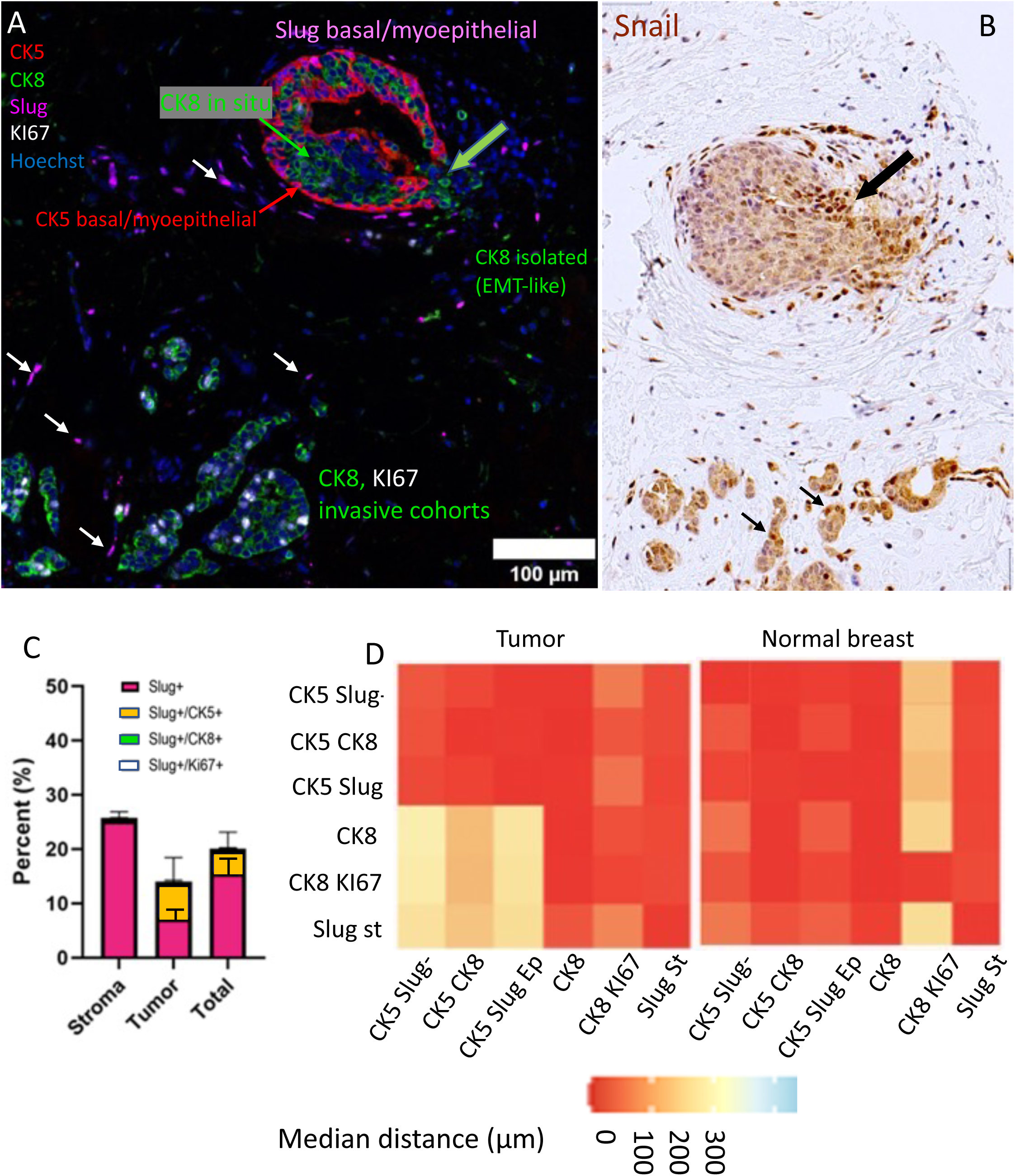
Spatial organization reflects cell differentiation and morphogenic events such as duct rupture. A. Multiplex analysis of duct rupture in a LumB breast carcinoma. CK8+ tumor cells are found in situ (green thin arrow), but also isolated and invading as cohorts, including most KI67+ cells. Myoepithelial CK5+ cells show a rift in the basal layer (fat arrow), providing passage to CK8+ isolated tumor cells. Slug is mostly expressed by CK5+ basal/myoepithelial cells, and isolated stroma cells (small white arrows). B. In an adjacent section, Snail is seen expressed preferentially by CK8+ tumor cells migrating out from the duct (arrow). C. Quantification using QuPath software confirms that Slug is about exclusively expressed by stroma and CK5+ cells. D. Vicinity study using QuPath software show the regrouping of CK5+ cells with CK5+CK8+ cells. It also confirms that KI67+ proliferating cells tend to be isolated, as well as stroma Slug+ cells in the tumor microenvironment, when the distance between CK5+ and CK8+ drops in organized ducts from normal breast.

### Cellular topology by cell proximity analysis

Using cell proximity analysis, we investigated the positioning of Slug-positive cells within tumor subpopulations (Fig. 6D). We found in all 3 LumB samples that CK5+Slug+ myoepithelial cells were always located in a close proximity to CK5+ clusters, including the hybrid CK5+CK8+ subset, suggesting a specific microenvironment including the less differentiated phenotype CK5+CK8+. Conversely, CK8+ cells were not at close proximity from CK5+CK8+ cells and formed separate clusters. CK8+KI67+ cells were scattered within these clusters, not at close proximity from each other, meaning there were mostly isolated among CK8+ cells.

We also looked at the location of the Slug+ stroma cells. Interestingly, we found these fibroblast-like cells to be located at a relative proximity from the CK8+KI67+cells, suggesting potential stroma-cancer cells interactions (Fig. 6D).

### Clinical impact: Slug expression pattern (H-scores) fail to show any significant link to the pathology prognostic when Snail expression pattern (H-scores) reflects the pathology prognostic in most cases but also uncover unexpected links

We first examined potential links between expression patterns of Slug and the prognosis of breast cancer patients from our cohort using the multivariable Cox model adjusted for the classical known risk factors. No association could be found between the prognostic and tumoral expression levels (HR_iDFS_ = 0.89, 95%CI [0.61; 1.29], p = 0.5457; HR_DDFS_ = 0.95, 95%CI [0.64; 1.41], p = 0.7890; HR_OS_ = 1.17, 95%CI [0.71; 1.93], p = 0.5448) or stromal expression levels (HR_iDFS_ = 1.15, 95%CI [0.80; 1.65], p = 0.4641; HR_DDFS_ = 1.08, 95%CI [0.73; 1.59], p = 0.6979; HR_OS_ = 1.13, 95%CI [0.69; 1.85], p = 0.6317).

We also looked at Snail expression pattern. The increase of the Snail tumoral expression was associated to an increase and an improvement of the prognostic (HR_iDFS_ = 0.81, 95%CI [0.67; 0.99], p = 0.0359; HR_DDFS_ = 0.79, 95%CI [0.64; 0.97], p = 0.0249; HR_OS_ = 0.76, 95%CI [0.57; 1.01], p = 0.0625). Using this methodology, the stromal expression did not seem to be associated with the patient prognostic (HR_iDFS_ = 1.03, 95%CI [0.84; 1.25], p = 0.8094; HR_DDFS_ = 1.03, 95%CI [0.83; 1.28], p = 0.7944; HR_OS_ = 1.00, 95%CI [0.74; 1.35], p = 0.9909).

Adjusting the statistical model for both tumoral and stromal expression, we observed a suppressor effect (Table 3), enhancing the association of the Snail tumoral expression with a better prognostic, and revealing the opposite effect of the increase of the stromal expression which seemed to be associated to a worse prognostic. As the proportional hazard did not hold for this association of the stromal expression with the iDFS, we introduced an interaction with time, showing that the risk increased with the value of the stromal expression and the time (HR_iDFS_(t) = 1.08, 95%CI [1.03; 1.13]), suggesting an association with of late invasive disease. Using spline did not reveal particular trend to define a time cut-off, suggesting that the risk increased linearly with the time from the surgery (Fig. 7, Table 3).

**Figure 7.**
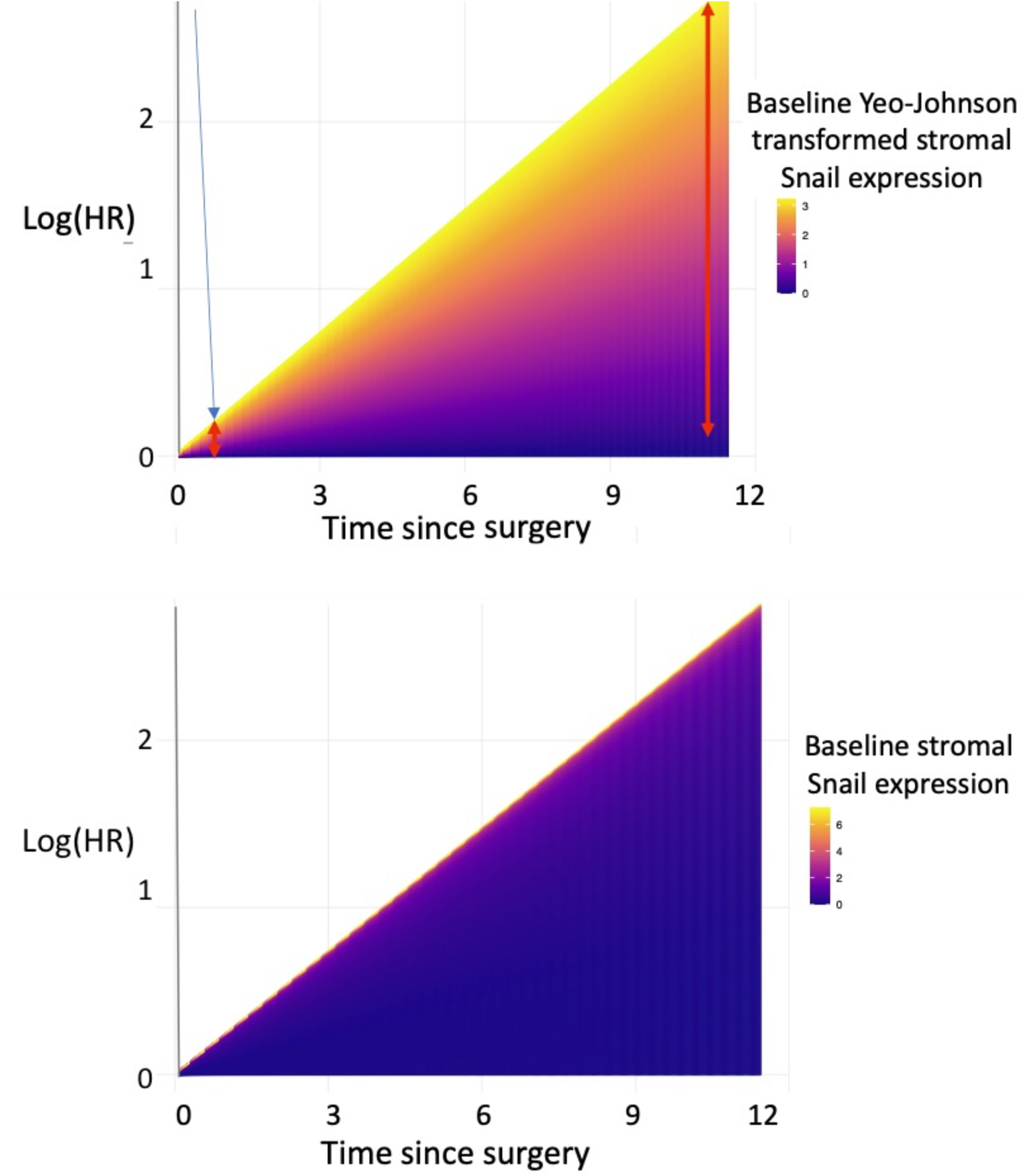
Time-varying association between the iDFS and the baseline stromal Snail expression. (Left: Yeo-Johnson transformed expression; right: raw expression). The worsen of the prognostic (Log (HR) with the increase of the baseline stromal Snail expression is low at baseline, but increase with the time from surgery.

**Table 3:** Association between the patient prognostic and the expression of the tumoral and stromal Snail expression. All models were adjusted for the classical risk factors (age at surgery, tumor size, tumor grade, luminal status, number of positive nodes, and the Ki67).

| Clinical endpoint | Model adjustment for<br>tumoral and stromal<br>expression | HR of the Snail<br>tumoral<br>expression<br>[95%CI] | p-value | HR of the Snail<br>stromal<br>expression<br>[95%CI] | p-value |
| --- | --- | --- | --- | --- | --- |
| iDFS<br>(n events=107) | Adjusted for only one | 0.81 [0.67; 0.99] | 0.0359 | 1.03 [0.84; 1.25] | 0.8094 |
|  | Adjusted for both | 0.69 [0.54; 0.89] | 0.0049 | 1.30 [1.00; 1.68] | 0.0487 |
|  | Adjusted for both with<br>time/stromal<br>expression interaction | 0.67 [0.53; 0.85] | 0.0009 | 1.08 [1.03; 1.13] | 0.0020 |
| DDFS<br>(n events = 93) | Alone | 0.79 [0.64; 0.97] | 0.0249 | 1.03 [0.83; 1.28] | 0.7944 |
|  | Adjusted for both | 0.66 [0.50; 0.86] | 0.0027 | 1.35 [1.02; 1.78] | 0.0348 |
| OS<br>(n events = 51) | Alone | 0.76 [0.57; 1.01] | 0.0625 | 1.00 [0.74; 1.35] | 0.9909 |
|  | Adjusted for both | 0.63 [0.43; 0.92] | 0.0181 | 1.36 [0.92; 2.00] | 0.1224 |

As exploratory analyses, we introduced interactions between different expressions and the luminal/hormonal/Her2 status in order to identify potential subgroups. Only an interaction between the stromal Snail expression and the Her2 status was found, still indicating a better prognostic with the increase of the tumor expression (HR_iDFS_ = 0.65, 95%CI [0.52; 0.83], p = 0.0005; HR_DDFS_ = 0.62, 95%CI [0.48; 0.81], p = 0.0003; HR_OS_ = 0.61, 95%CI [0.42; 0.88], p = 0.0076), and a worsen prognostic with the stromal expression and time (HR_iDFS_(t) = 1.06, 95%CI [1.01; 1.11], p = 0.0192; HR_DDFS_ = 1.05, 95%CI [1.00; 1.11], p = 0.0674; HR_OS_ = 1.03, 95%CI [0.96; 1.11], p = 0.3482), but this last was specifically worsened for the Her2 positive patients (HR_iDFS_ = 2.33, 95%CI [1.24; 4.38], p = 0.0087; HR_DDFS_ = 2.89, 95%CI [1.50; 5.60], p = 0.0016; HR_OS_ = 9.45, 95%CI [2.44; 36.58], p = 0.0011).

## DISCUSSION

In this manuscript, we focused on two major EMTaTF, Snail and Slug, and identified for the first time by immunochemistry cell types expressing Slug and Snail protein in a significant cohort of more than 500 invasive breast carcinomas. Our results put some perspective on the previous pre-clinical studies on Slug and Snail clinical impact on tumor progression. Clinically, Slug expression in tumor cells was not found overall to be linked to clinical progression when Snail, unexpectedly, was linked to a better one. These results do not support previous studies based on Slug or Snail RNA expression level typically concluding on a positive link to tumor progression and metastasis. This probably reflects the fact that we based our conclusions on the protein level from distinct cell types, not a whole tumor RNA. It is well demonstrated that Slug or Snail overexpression can induce an EMT in various epithelial transformed or normal cell types (our work and others). However, our results show that the clinical outcome also depends on other pathways, potentially involving EMTaTF playing a distinct context-dependent role such as cell differentiation, proliferation regulation, immune and drug resistance with a specific impact on clinical progression. In any case, most of the tumor cells expressing Slug or Snail did not express an individualized phenotype, preserving at least some clearly epithelial features, such as cytokeratin expression and overall cohesiveness with neighbor cells. One general conclusion of this study is that we could not link Snail or Slug expression with extensive EMT in our extensive panel of breast carcinoma, without precluding intermediate and partial EMT stages, supported by the visible lack of apical-basal polarity in most carcinoma cells (Figs 2 and 3).

We described earlier, with other laboratories, the expression of Slug in a minor subpopulation of myoepithelial cells in healthy breast samples, associated with proliferation (Weinberg, Nassour et al 2012). In fact, Slug basal expression is involved in cell proliferation and differentiation in most pluri-stratified epithelia, including basal keratinocytes (Savagner et al 2005). Here, we find Slug to be over-expressed by most normal-looking myoepithelial cells in tumors such as luminal/ER+ and HER2 carcinoma. These CK5+Slug+ myoepithelial cells displayed a normal phenotype but remarkably were not proliferating (KI67-), suggesting a specific activation state apparently linked to the tumor microenvironment. More Slug was found to be expressed in LumA tumor cells (grade 1) as compared to LumB (grade 1), apparently in link with a larger extent of normal looking tubule structures. In TN carcinoma, Slug was significantly expressed by invasive groups of tumor cells, but only in a minority (25%) of our samples, without any visible tropism for specific tumor areas. Cells expressing Slug were typically isolated among Slug- tumor cells in most cases.

Using vicinity measurements, we characterized two types of cell clusters linked to Slug pattern of expression. Among other tumor cells, Slug+ cells were located closer from basal CK5+ cells, including Slug- cells, but also flanked a hybrid CK5+CK8+ subpopulation, suggesting a filiation between these subpopulations. Conversely, CK5-CK8+ luminal cells formed more cohesive and homogeneous groups (Fig. 7). The emergence of a double positive CK5+CK8+ subpopulation expressing a hybrid phenotype is not uncommon in breast carcinoma. It is typically linked to a transition phase to a more aggressive phenotype for the tumor, as described in two breast cancer mouse models using microarrays and single cell analysis (Rädler et al., 2021; Sinha et al., 2021).

The second cluster that we observed was unexpected, we found the Slug+CK- stroma cells involved a higher percentage of stroma cells than in normal mammary gland. We found that these cells were located relatively close from the proliferative luminal CK8+KI67+ tumor cells that we described by multiplex analysis, dispersed among CK8+ tumor cells. Added to the correlation that we found between tumoral KI67 expression level and stromal Slug+ cells, it suggests that the slug expression by carcinoma-activated fibroblasts is linked to the tumor proliferation level, potentially involving a shared pathway or some other mechanism in Luminal tumors. This may contribute to a clinical progression linked to higher Slug expression in stroma cells, but we could not observe it directly, confirming that proliferation is just one of the factors contributing to tumor progression. Nevertheless, this hypothesis is also supported by our finding that stromal Slug expression level is twice higher in higher grade (Grade 2-3) luminal B tumors as compared to lower grade (grade 1) tumor (Fig.1G).

In contrast, Snail was frequently expressed in transformed cells in all invasive breast carcinomas subtypes. Expression pattern was heterogeneous and some stroma cells, particularly endothelial cells, as confirmed by CD31 co-staining, were also found to express Snail in most tumor subtypes, suggesting a specific activation state possibly linked to neovascularization mechanisms, involving endothelial cell plasticity and invasiveness. We also found Slug to be expressed in endothelial cells, but only in rare cases in our sampling.

The main surprise in our study was the link between tumor Snail protein expression and a better prognosis overall in our sampling (Table 4). We found Snail to be expressed in a limited subpopulation in all tumor subtypes, involving invasive areas and in situ component, even in the less aggressive forms. Staining was very heterogeneous between tumor cells.

**Table 4:** Summarizing the clinical impact of Snail and Slug based on their cell expression pattern.

|  | Cellular Localization (nuclei) | Functional/Clinical impact |
| --- | --- | --- |
| **Slug** | Epithelial<br>(tumor or normal cells) | No significant link |
|  | Stromal cells | Positive correlation between stromal Slug and KI67 expression.<br>No significant impact on prognosis. |
| **Snail** | Epithelial<br>(tumor or normal cells) | Positive link between Snail expression and prognosis improvement (all types) |
|  | Stromal cells | Time-linked worsening prognostic<br>(potential link to relapse, metastasis) |

The stromal Snail expression seemed to have an opposite effect : an increase worsened the prognosis. Interestingly, this worsening was more important for the patients with the HER2 subtype, a subtype described to include a particularly abundant tumor-linked endothelial network (Konecny et al. 2004), suggesting neo-vasculature including Snail overexpression could play a worsening role in clinical progression in HER2 subtype and potentially other subtypes. This hypothesis is supported by mouse model MMTV-PyMT in which Snail depletion in endothelial cells is linked to a better survival rate (Cabrerizo-Granados et al 2021). Overall, it appears that Snail could be involved in distinct pathways beyond EMT promotion, linked to an impact on cell differentiation inducing a less aggressive phenotype in some tumor microenvironments.

Slug and Snail expression pattern along tumor histological subtypes followed an expectable profile: Slug was overexpressed in tubular subtype, characterized by the relative abundance of tubules and myoepithelial cells. Slug and Snail were also overexpressed in our few samples of metaplastic type (Fig.1F), a subtype characterized by an impaired differentiation evoking a partial but visible EMT. In contrast, the lowest Slug expression level was detected in papillary subtype, in which tumor cells display a reversed polarity with an attenuated myoepithelial cells layer. These localizations supported the link between Slug expression and differentiation pattern, rather than EMT occurrence.

This work is now being completed by an immunolocalization of other EMT associated transcription factors Twist and Zeb, to complete our investigation on the localization pattern and impact of the main EMT associated transcription factors during breast cancer progression. This will provide a more comprehensive view on the roles of these pathways during cancer progression.

## ACKNOWLEDGEMENTS.

This work is supported by the Ligue Nationale contre le cancer (2021 grant: R21078LL), by a grant from Gustave Roussy (Taxe apprentissage) and by the Prism project funded by the Agence Nationale de la Recherche under grant number ANR-18-IBHU-0002. F.D. received a PhD fellowship from Prism-ANR and is laureate from the “Ligue Nationale contre le cancer” 4th year thesis fellowship. We are very thankful to Dr. Barbara Pistilli for the helpful discussions and support for accessing precious TMA samples.

## Authors contribution

F.D. contributed directly to the acquisition, analysis, and interpretation of data. F.D. also contributed to the writing reviewing and editing. L.M. contributed to the reviewing and editing. D.D. as a Gustave Roussy staff biostatistician covered the intricate statistical analysis. B.V. as a Gustave Roussy clinical oncologist contributed to the analysis and interpretation of data. N.J. as a Gustave Roussy clinical pathologist, overlooked analysis and interpretation of data; P.S. mostly contributed to the conception, design of the work, supervision and writing. We thank the PFEP from Gustave Roussy (M. Polrot, Plateforme d’Evaluation Préclinique), for the support with sample processing.

## FIGURE LEGENDS

**Supp Figure 1.**
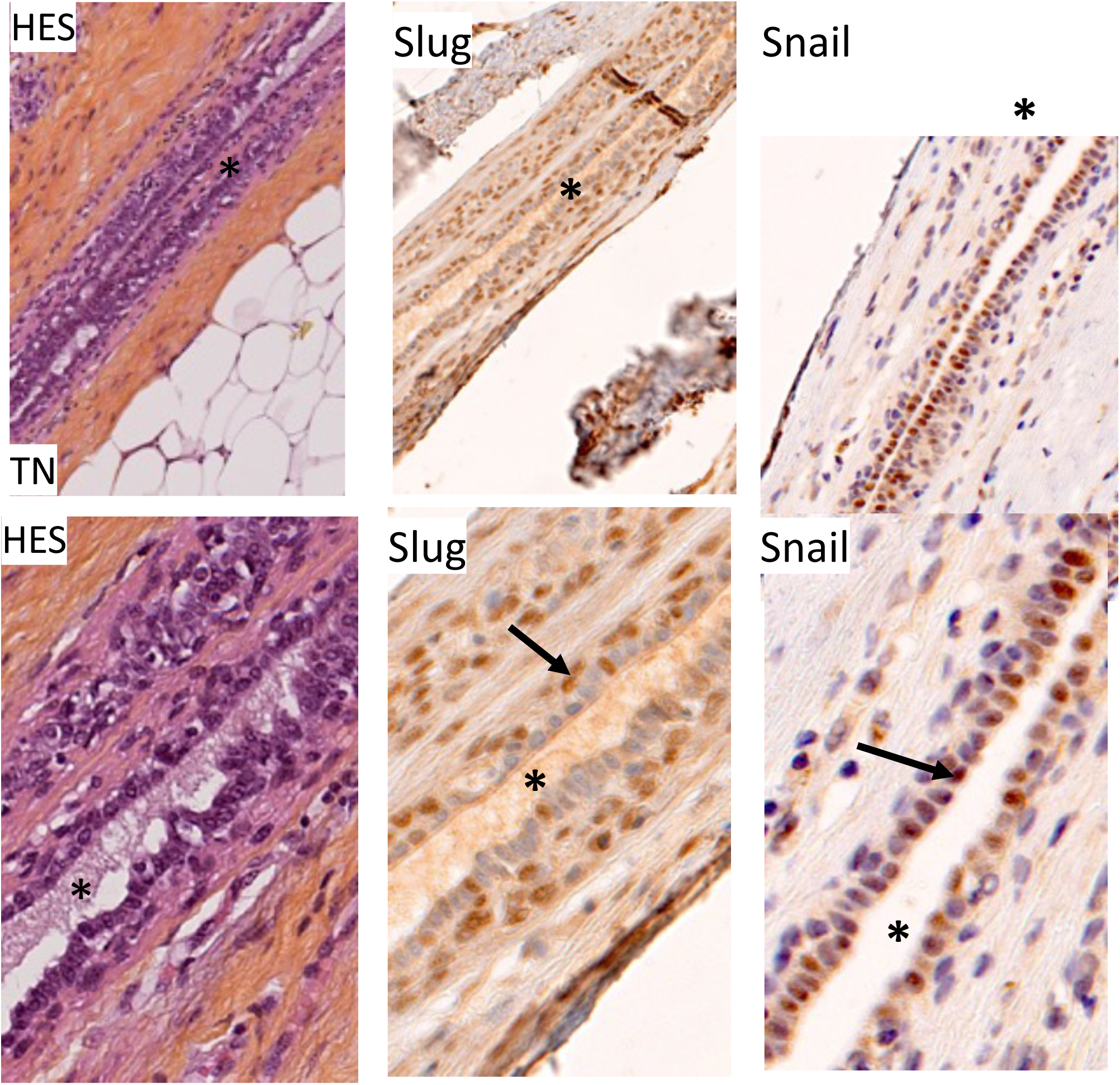
Rare expression of Snail in normal-looking luminal cells inside a TN breast carcinoma. Lower panel include higher magnification from upper panel regions. HES staining shows a tubule section including some mucus in the lumen (*). Slug is expressed in basal cells, as seen in other figures (Center panel, arrow). Snail is expressed in normal-looking cells (right panel, arrow).

